# Arabidopsis Lunapark proteins are involved in ER cisternae formation

**DOI:** 10.1101/256743

**Authors:** Verena Kriechbaumer, Emily Breeze, Charlotte Pain, Frances Tolmie, Lorenzo Frigerio, Chris Hawes

## Abstract

The plant endoplasmic reticulum (ER) is crucial to the maintenance of cellular homeostasis. The ER consists of a dynamic and continuously remodelling network of tubules and cisternae. Several conserved membrane proteins have been implicated in formation and maintenance of the ER network in plants, such as RHD3 and the reticulon family of proteins.

Despite the recent work in mammalian and yeast cells, the detailed molecular mechanisms of ER network organisation in plants still remain largely unknown. Recently novel ER network-shaping proteins called Lunapark have been identified in yeast and mammalian cells.

Here we identify two arabidopsis LNP homologues and investigate their subcellular localisation via confocal microscopy and potential function in shaping the ER network using protein-protein interaction assays and mutant analysis.

We show that AtLNP1 overexpression in tobacco leaf epidermal cells mainly labels the three-way junctions (trivia) of the ER network whereas AtLNP2 labels the whole ER. Overexpression of LNP proteins results in an increased abundance of ER cisternae and an *lnp1lnp2* amiRNA line displays a less structured ER network.

Thus, we hypothesize that AtLNP1 and AtLNP2 are involved in determining the dynamic morphology of the plant ER, possibly by regulating the formation of ER cisternae.

## Introduction

As the first biosynthetic organelle in the plant secretory pathway, the endoplasmic reticulum (ER) underpins the production, folding and quality control of proteins (Brandizzi *et al.*, 2003; Hawes *et al.*, 2015), as well as lipid biosynthesis (Wallis and Browse, 2010). In addition, it also has many other functions such as calcium homeostasis (Hong *et al.*, 1999), oil and protein body formation (Huang, 1996; Herman, 2008), rubber particle formation (Brown *et al.*, 2017) and auxin regulation (Friml and Jones, 2010; Kriechbaumer *et al.*, 2015).

In plant cells, the ER consists of a dynamic network of cisternae (sheets) and, more predominantly, tubules which extend throughout the cytoplasm and across cellular boundaries, with intimate connections to other organelles such as the Golgi, the plasma membrane, the outer membrane of the nuclear envelope and the cytoskeleton (Goyal and Blackstone, 2013; Hawes *et al.*, 2015; Stefano and Brandizzi, 2017). Hence, the ER is crucial to the maintenance of cellular homeostasis. The ER network is continually remodelling, presumably in response to differing cellular demands, with the formation of new three-way junctions and polygons via tubule extension and fusion events, balanced with polygon loss from tubule sliding and ring closure (Griffing, 2010). Several conserved membrane proteins have been implicated in both the formation and maintenance of the ER network in plants, notably the GTPase ROOT HAIR DEFECTIVE3 (RHD3) and the Reticulon (RTN) family of proteins. RHD3, orthologous to mammalian atlastins (ATL) and yeast Sey1p, may mediate membrane fusion and the formation of three-way junctions (Zhang *et al.*, 2013; Chen *et al.*, 2011). These three-way junctions of the ER, which we propose to rename ER ‘trivia’ (from the Latin *trivium*, place where three roads meet), consist of small triangular sheets with concave edges (Shemesh *et al.*, 2014). The fusogenic action of RHD3 is then complemented by the curvature-generating and/or stabilizing RTN proteins, which preferentially localise to ER tubules and the curved edges of cisternae. We recently demonstrated that an arabidopsis RTN (RTN13) relies on a conserved amphipathic helix to induce membrane curvature *in vivo* (Breeze *et al.*, 2016). Indeed, purified yeast and mammalian orthologs of these two groups of proteins (Sey1p or ATL with RTNs) are sufficient to reconstitute a dynamic tubular ER network in proteoliposomes in the presence of GTP (Powers *et al.*, 2017).

A third class of conserved ER network-shaping proteins called Lunapark (Lnp1p [yeast] and mLnp1 [mammals]) has additionally been identified in yeast and mammalian cells. Lunapark (LNP) proteins are characterised by the presence of two N-terminal transmembrane domains (TMDs) and an atypical Cys4 type C-terminal zinc finger motif which, in yeast, mediates homodimerization and is required for LNP function (Casey *et al.*, 2015; Wang *et al.*, 2016). Immediately adjacent to the zinc finger lies the amino acid sequence LNPARK or a variant thereof. Mammalian Lnp1 is dependent on N-myristoylation for its localisation to ER junctions and morphogenic activity. The absence of this motif in yeast Lnp1p may indicate that LNP proteins in higher organisms have evolved an additional level of functionality and/or regulation via post-translational lipid modifications (Moriya *et al.*, 2013; Turnbull *et al.*, 2017, Wang *et al.*, 2016).

LNP proteins preferentially localise to ER trivia *in vivo* and, although they are not required for ER network formation, it has been suggested that LNP acts to stabilise these intersections, potentially acting as temporary scaffold in nascent junctions (Chen *et al.*, 2012; Chen *et al.*, 2015; Wang *et al.*, 2016). Immunofluorescence staining of mammalian COS-7 cells with an anti-mLnp1 antibody showed that only around half of all trivia contained mLnp1 but that mLnp1 acquisition was associated with enhanced junction survival probability and reduced junction mobility and ring closure (Chen *et al.*, 2015).

Lnp1p has been reported to work synergistically with RTNs and Yop1 (an additional curvature stabilizing protein found in yeast, orthologous to mammalian REEPs), but in antagonism to Sey1p to maintain the cortical ER network in yeast cells (Chen *et al.*, 2012). Indeed, in *sey1*Δ yeast mutants lacking functional Sey1p, Lnp1p is expressed in cisternae throughout the ER network, including the nuclear envelope, suggesting that in the wildtype Sey1 acts upstream of Lnp1 restricting Lnp1 to ER tubule junctions (Chen *et al.*, 2012). Co-immunoprecipitation of epitope-tagged Lnp1p with RTN1p, Sey1p or Yop1p proteins, showed that Lnp1p was capable of interacting with all three yeast ER morphogens *in vivo*, and that, moreover, the interaction of Lnp1p with Rtn1p is negatively regulated by Sey1p (Chen *et al.*, 2012). However, more recently Wang *et al.* (2016) reported that in mammalian cells stably expressing both Lnp-mCherry and GFP-ATL-3, ATL was in close proximity to Lnp-mCherry in the peripheral ER network but, crucially, the two proteins did not precisely co-localise and no interaction was found in reciprocal pull-down experiments.

Despite the recent work in mammalian and yeast cells, the detailed molecular mechanisms of ER network organisation in plants still remain largely unknown. Here we identify two arabidopsis LNP homologues and show that their overexpression (as fluorescent protein fusions) in tobacco leaf epidermal cells results in the proteins labelling the ER and accumulating at the trivia of the ER network. Furthermore, overexpression of LNP proteins results in an increased abundance of ER cisternae in the ER network. Thus, we hypothesize that AtLNP1 and AtLNP2 are involved in determining the dynamic morphology of the plant ER, possibly by regulating the formation of ER cisternae at three-way junctions.

## Materials and Methods

### Cloning of expression plasmids

Primers were obtained from Eurofins Genomics. Q5 high-fidelity DNA polymerase (New England Biolabs) was used for all PCR reactions. Genes of interest were cloned with a C-terminal GFP fusion under a ubiquitin-10 promoter (*P*_*UBQ10*_) (Grefen *et al.*, 2010) using Gateway technology (Invitrogen).

### Preparation of *lnp1lnp2* amiRNA line

Candidate amiRNA sequences specific to both *LNP1* and *LNP2* coding regions were identified using the Web MicroRNA Designer (WMD) platform (http://wmd3.weigelworld.org) (Ossowski *et al.*, 2008; Schwab *et al.*, 2006). Two amiRNA sequences were selected (targeted against the conserved zinc finger domain [amiRNA1] or the region immediately downstream of TMD2 [amiRNA2]) and cloned into the naturally occurring arabidopsis miR319a replacing the target specific sequence using a series of overlapping PCRs (as described by Ossowski *et al.*, 2008) and with the addition of Gateway-compatible attB sites. The purified attB-amiRNA precursors were subsequently used to generate Entry (pDONRZeo) and 35S destination (pB7WG2) clones. The 35S constructs were transformed by heat-shock into *Agrobacterium tumefaciens* strain GV3101 and stable homozygous arabidopsis lines created via the floral dip procedure (Clough and Bent, 1998). RNA was extracted from dry seeds of the amiRNA1- and amiRNA2-containing lines and Col-0 as described by (Meng and Feldman, 2010) and first-strand cDNA synthesized using ReadyScript (Sigma Aldrich) according to the manufacturer’s instructions. Semi-quantitative RT-PCR was performed using primers specific to LNP1 and LNP2, together with At4g34270 and At4g12590 (seed-specific housekeeping reference genes 1 and 2, respectively (Dekkers *et al.*, 2012)) as loading controls, and expression levels in the amiRNA mutants compared to that of wildtype, Col-0. The amiRNA2-containing line was found to have reduced expression of *LNP1* but comparable expression of *LNP2* to that detected in wild-type and was subsequently referred to as *lnp1*; whereas the amiRNA1-containing line had markedly reduced expression levels of both *LNP1* and *LNP2* and was therefore denoted as *lnp1lnp2*.

### Plant material and transient expression in tobacco leaves

For Agrobacterium-mediated transient expression, 5-week-old tobacco (*Nicotiana tabacum* SR1 cv Petit Havana) plants grown in the greenhouse were used. Transient expression was carried out according to Sparkes *et al.* (2006). In brief, each construct was introduced into Agrobacterium strain GV3101 by heat shock. Transformants were inoculated into 3 ml of YEB medium (per litre: 5 g of beef extract, 1 g of yeast extract, 5 g of sucrose and 0.5 g of MgSO_4_·7H_2_O) with 50 μg ml^−1^ spectinomycin and 25 μg ml^−1^ rifampicin. After overnight shaking at 25°C, 1 ml of the bacterial culture was pelleted by centrifugation at 2200 *g* for 5 min at room temperature. The pellet was washed twice with 1 ml of infiltration buffer (50 mM MES, 2 mM Na_3_PO_4_·12H_2_O, 0.1 mM acetosyringone and 5 mg ml^−1^ glucose) and then resuspended in 1 ml of infiltration buffer. The bacterial suspension was diluted to a final OD_600_ of 0.05 (0.01-0.3 in the OD_600_ serial dilution series) and carefully pressed through the stomata on the lower epidermal surface using a 1 ml syringe. Infiltrated plants then were returned to glasshouse conditions for 48 h prior to imaging.

### Confocal microscopy

Images were taken using a Zeiss 880 laser scanning confocal microscope with ×100/1.4 NA oil immersion objective. For imaging of the GFP/RFP combinations, samples were excited using 488 and 543 nm laser lines in multi-track mode with line switching. Signals were collected using either standard GaAsP detectors or the high resolution Airyscan detector. Images were edited using the ZEN image browser.

### Lipid dye Rhodamine B hexyl ester

Staining the ER with Rhodamine B hexyl ester was carried out according to Hawes *et al.* (2018). Rhodamine B hexyl ester solution was prepared as a 1 mM stock solution in DMSO and a 1 μM working solution in distilled water (DW). Whole arabidopsis seedlings 7-10 days after germination were transferred to Eppendorf tubes containing 1 μM Rhodamine B hexyl ester. Seedlings were incubated for 15 minutes in the dye and washed in DW. Samples were imaged with a 514 nm argon ion laser emission detected using 470-500 nm and 560-615 nm band-pass filters.

### FRET-FLIM Data Acquisition

Constructs were transiently expressed in tobacco leaf epidermal cells as described above. Leaf discs were excised and the GFP and mRFP expression levels in the plant within the region of interest were confirmed using a Nikon EC2 confocal microscope at 488 and 543 nm, respectively. FRET-FLIM data capture was performed according to Kriechbaumer *et al.* (2015) using a two-photon microscope at the Central Laser Facility of the Rutherford Appleton Laboratory. A two-photon microscope built around a Nikon TE2000-U inverted microscope was used with a modified Nikon EC2 confocal scanning microscope to allow for multiphoton FLIM (Schoberer and Botchway, 2014). At least three nuclei from at least two independent biological samples per protein-protein combination were analysed, and the average of the ranges was taken.

### Quantification of ER polygonal regions in WT and mutants

Images acquired with a Zeiss 880 confocal microscope with Airyscan (see above) were analysed with ImageJ ‘Analyse particles’. In preparation for the analysis images were processed as follows: a smoothing function was applied in order to reduce noise; images were then binarised using Otsu’s method (Otsu, 1979); a closing function was applied to ensure high connectivity of the network; the image LUT was inverted as this was required for next step; ImageJ’s ‘Analyse particles’ package was then used to quantify the polygonal region size. Only Polygonal regions completely enclosed by the network were considered for the analysis. As the data is not normal, the Wilcoxon rank sum test was applied. p-value < 2.2^e-16^ were calculated for three biological replicas with at least 12 technical repeats each.

### Persistency analysis

To analyse persistency of LNP1-labelled punctae and cisternae, movies were taken using the Zeiss 880 confocal with Airyscan. RFP-HDEL was used as an ER luminal marker. The ImageJ plugin ‘temporal color-code’ was applied to colour-code movement over time. In this case in 3 image frames at 0, 30 and 60 s. White areas indicate points of persistency throughout in the composite image, red areas indicate highest mobility.

## Results

### Identification of two Lunapark orthologues in arabidopsis

The protein sequences of yeast Lnp1p and human Lnp were queried against the *Arabidopsis thaliana* proteome using BLASTP. This analysis identified two Lunapark orthologues, At2g24330 and At4g31080 (subsequently named AtLNP1 and AtLNP2, respectively), both with ~24% amino acid identity to the queried sequences across the entire length of the protein. AtLNP1 and 2 themselves share 67% amino acid identity. According to large scale microarray data in the eFP browser (Winter *et al.*, 2007) both At*LNP1* and At*LNP2* are transcribed ubiquitously but At*LNP2* has increased transcription levels in pollen (Supplementary Figure S1). Further comparison of the identified AtLNP protein sequences with their mammalian and yeast orthologues revealed the presence of several conserved features, notably two TMDs towards the N-termini and a zinc finger motif immediately adjacent to the LNPARK motif (LNKPKH in AtLNP1 and 2) (Figure 1). However, unlike the human (and mouse) Lnp proteins, AtLNP1 and 2 do not contain a Pro-rich domain upstream of the zinc finger motif (Figure 1), although the functional significance of this domain in mammalian Lnp proteins is currently unknown. The Lunapark amino acid motif is conserved throughout the *Viridiplantae* kingdom (green algae and land plants), although some variation in specific residues exists between species. Multiple sequence alignment of 161 identified Lunapark homologs from 55 species within the green plant lineage using the PLAZA 4.0 online platform (Proost *et al.*, 2009) identified a consensus sequence motif of [A/G][L/K][N/E]XPK (Supplementary Figure S2).

**Figure 1:**
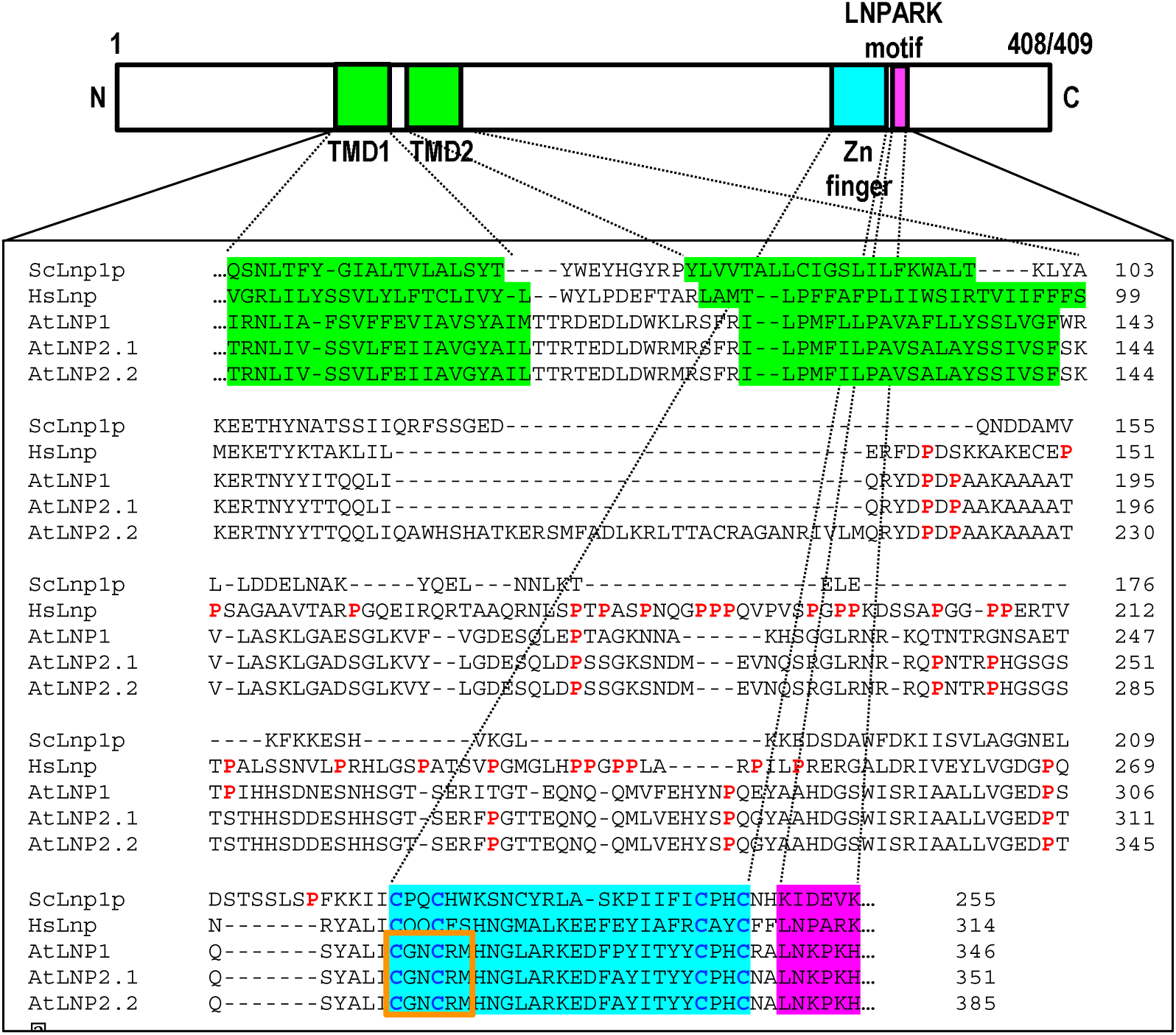
AtLNP1 and AtLNP2 are orthologous to Lunapark proteins in other organisms. Two putative Lunapark (LNP) proteins were identified in *Arabidopsis thaliana*, with sequence homology and conserved motifs (two transmembrane domains, TMDs; zinc finger, and LNPARK motif [LNKPKH in arabidopsis]) with previously annotated *Homo sapiens* Lnp (HsLnp) and *Saccharomyces cerevisiae* Lnp1p (ScLnp1p) proteins. Note: Unlike yeast and arabidopsis LNP proteins, HsLnp contains an additional Pro-rich region (Pro residues shown in red). The orange boxed sequence shows the conserved region targeted by an artificial microRNA in *lnp1lnp2* loss-of-function lines. AtLNP1 and AtLNP2.1 are 408 and 409 amino acid residues in length, respectively.

The mammalian Lnp protein features an N-terminal myristoylation site and ER network changes induced by overexpression of Lunapark proteins were significantly inhibited by the mutation of protein N-myristoylation which rendered this motif non-functional (Moriya *et al.*, 2014). Yeast Lnp1p protein in contrast does not feature this motif (Moriya *et al.*, 2014). Arabidopsis LNP proteins similarly do not possess an N-terminal glycine and therefore are predicted to lack a myristoylation site as shown for the mammalian Lnp protein (Moriya *et al.*, 2014).

AtLNP2 is annotated as having two possible splice variants (AtLNP2.1 and AtLNP2.2) with AtLNP2.2 containing an additional 34 residues downstream of the TMDs. However, this region is not well conserved in LNP proteins from other organisms and moreover, publicly available RNASeq data (Araport) obtained from a range of tissues show no detectable sequence reads for this region. Hence, all subsequent analysis was performed on splice variant AtLNP2.1 (subsequently referred to as AtLNP2).

### AtLNP1 and AtLNP2 localise to different substructures within the ER network

To determine the subcellular location of AtLNP1 and AtLNP2, the full-length protein sequences were fused at the C-terminus to green fluorescent protein (GFP) under the control of the ubiquitin-10 promoter (*P*_*UBQ10*_) (Grefen *et al.*, 2010). Transient expression of both constructs in *Nicotiana tabacum* leaf epidermal cells alongside the ER luminal marker RFP-HDEL, showed specific subcellular localisation of AtLNP1 to the trivia regions of the ER network and to ER cisternae, but, notably, there was no detectable labelling of the ER tubules (Figure 2A). In contrast, whilst AtLNP2 was similarly observed to localise to ER sheets and trivia, it was additionally present in the ER tubular network (Figure 2B).

**Figure 2:**
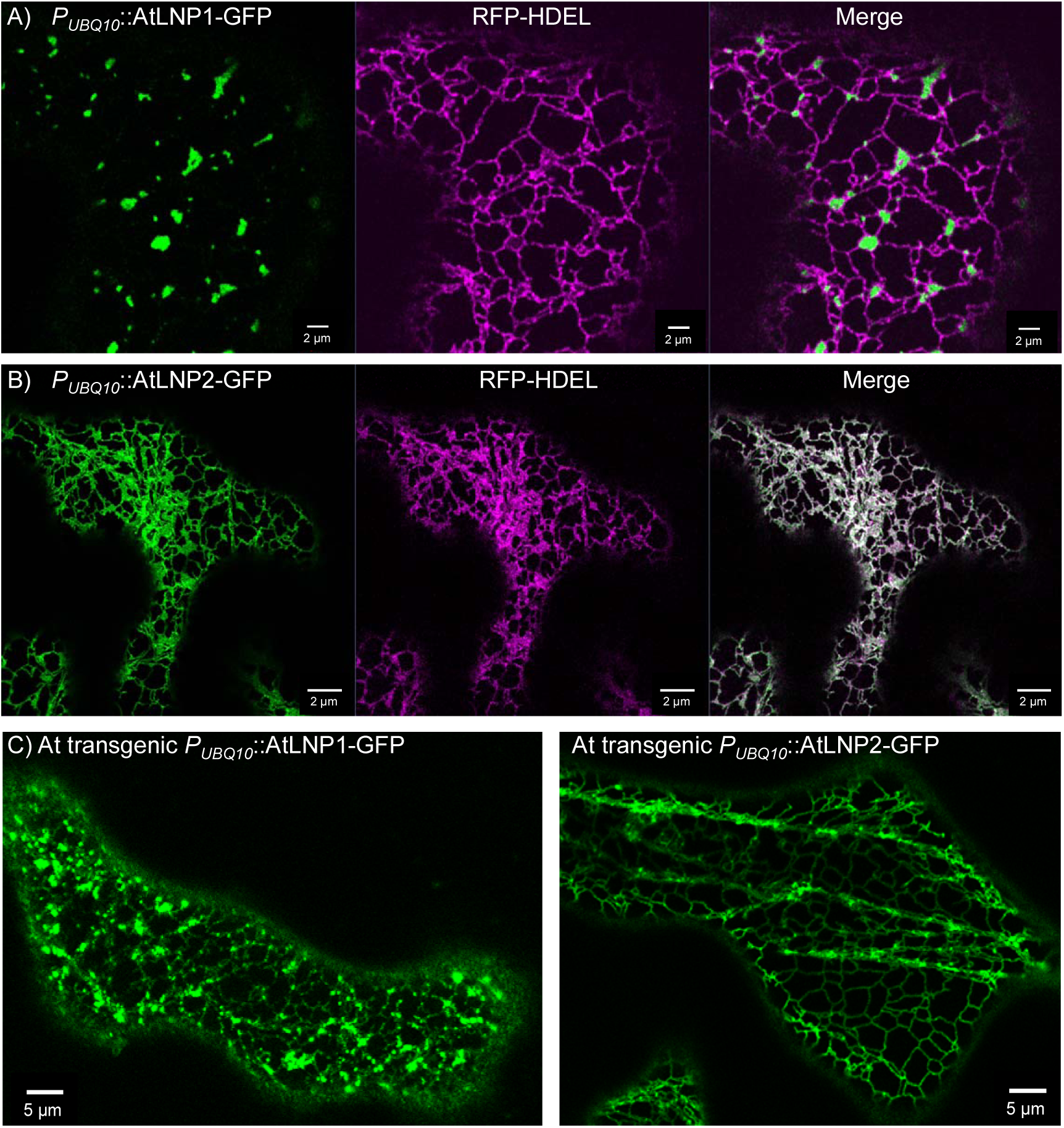
Localisation of AtLNP1 and AtLNP2 within the ER network. Transient co-expression of the ER luminal marker RFP-HDEL with (A) *P*_*UBQ10*_∷AtLNP1-GFP or (B) *P*_*UBQ10*_∷AtLNP2-GFP in *Nicotiana tabacum* leaf epidermal cells. Both AtLNP1 and AtLNP2 are localised to ER cisternae and trivia, but AtLNP2 additionally localises to ER tubules. C) Similar localisation patterns are shown for arabidopsis lines stably expressing *P*_*UBQ10*_∷AtLNP1-GFP or *P*_*UBQ10*_∷AtLNP2-GFP, respectively.

These two different location patterns were also found in arabidopsis plants stably expressing AtLNP1 or AtLNP2, respectively (Figure 2C).

### Expression of arabidopsis LNP proteins induces cisternae in a dose-dependent manner

The effect of protein expression dosage on the targeting of AtLNP1 and AtLNP2 to different ER structures was investigated by transient expression of *P*_*UBQ10*_∷AtLNP1-GFP and *P*_*UBQ10*_∷AtLNP2-GFP constructs across a range of Agrobacterium optical densities (ODs), as a proxy for differing protein expression levels within the agro-infiltrated leaf (Figure 3, Supplementary Figure S3). At higher doses of *P*_*UBQ10*_∷AtLNP1-GFP, AtLNP1-labelled sheets became increasingly prominent with the prevalence of ER cisternae at the highest OD of the *P*_*UBQ10*_∷AtLNP1-GFP construct (Figure 3, Supplementary Figure S3A). Moreover, at higher levels of AtLNP1 expression, some AtLNP1-labelled ER tubules were additionally observed (Figure 3, Supplementary Figure S3A).

**Figure 3:**
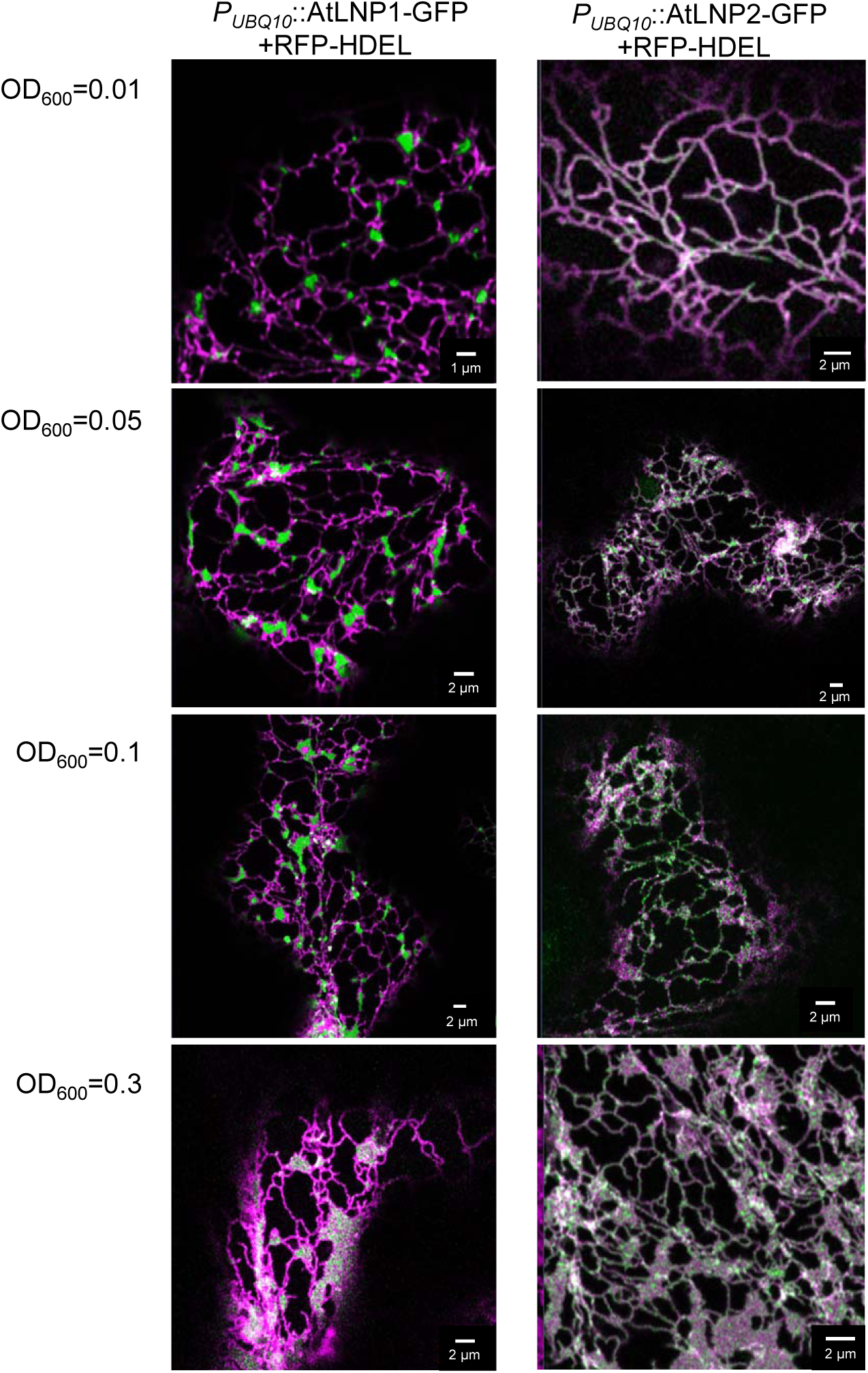
Increasing levels of protein expression affect the localisation of AtLNP1 and AtLNP2 within the ER network. Transient co-expression of *Agrobacterium tumefaciens* transformed with *P*_*UBQ10*_∷AtLNP1-GFP and *P*_*UBQ10*_∷AtLNP2-GFP, respectively, at increasing ODs (alongside the ER luminal marker, RFP-HDEL at a constant OD of 0.1) in *N. tabacum* leaf epidermal cells. At higher ODs an increasing formation of ER cisternae is observed for both constructs, and, for AtLNP1 additional labelling of ER tubules which is absent at lower ODs.

Protein dosage effects were also seen for AtLNP2, similarly achieved through agroinfiltration of *P*_*UBQ10*_∷AtLNP2-GFP at increasing ODs. At higher construct concentrations AtLNP2-GFP labelling of ER trivia became increasingly prevalent, together with a marked escalation of ER cisternae formation (Figure 3, Supplementary Figure S3B).

### An amiRNA *lnp1lnp2* loss-of-function mutant shows altered ER network morphology

Given the abundance of LNP proteins at trivia in the ER network and their ability to induce ER cisternae formation when expressed at high levels *in vivo*, we would hypothesise that downregulation of LNP could affect the number of trivia and therefore the overall appearance of the ER network. The effect on the ER network morphology in the absence of LNP proteins was therefore examined. Stable arabidopsis homozygous lines overexpressing GFP-HDEL and artificial microRNAs (amiRNA) designed to target either the conserved zinc finger region in both At*LNP1* and At*LNP2* (orange boxes in Figure 1) (*lnp1lnp2*) or a region immediately downstream of TMD2 (*lnp1*), were generated. Gene expression analysis showed expression levels of the relevant At*LNP* transcripts to be significantly reduced in these lines (Supplementary Figure S4). Loss of LNP had no noticeable effect on plant growth; unlike, for example, *rhd3* mutants which exhibit defects in plant development and are dwarfed (Zhang *et al.*, 2013).

The ER network in the stable homozygous *lnp1* amiRNA line was compared to Col-0 plants transformed with GFP-HDEL (Figure 4a). Cells from both lines were stained with the lipid dye Rhodamine B hexyl ester, which stains the ER but also mitochondria. No visible differences in ER morphology were observed between Col-0 and the *lnp1* mutant. In contrast, however, comparison of the ER network structure in cotyledons from Col-0 and the *lnp1lnp2* knockdown line, both stably transformed with GFP-HDEL, revealed obvious ER morphological differences (Figure 4b) whereby the polygons defined by the ER tubules appeared enlarged in *lnp1lnp2*. Subsequent quantitative *in silico* image analysis using ImageJ’s ‘Analyse particles’ showed that the average ER polygonal area in *lnp1lnp2* leaves was indeed significantly greater than in the corresponding GFP-HDEL controls (Figure 4c). In addition, this analysis revealed that the ER polygonal areas are less uniform and have a greater range of sizes than in the wildtype control (Figure 4c). Overall, this results in a less evenly structured ER network in the *lnp1lnp2* amiRNA leaves. The observed alterations in ER polygonal areas in the arabidopsis *lnp1lnp2* line here may be due to a loss of trivia and thus, cisternae originating from these junctions, or a reduction in the trivia stability resulting in more irregular polygonal areas.

**Figure 4.**
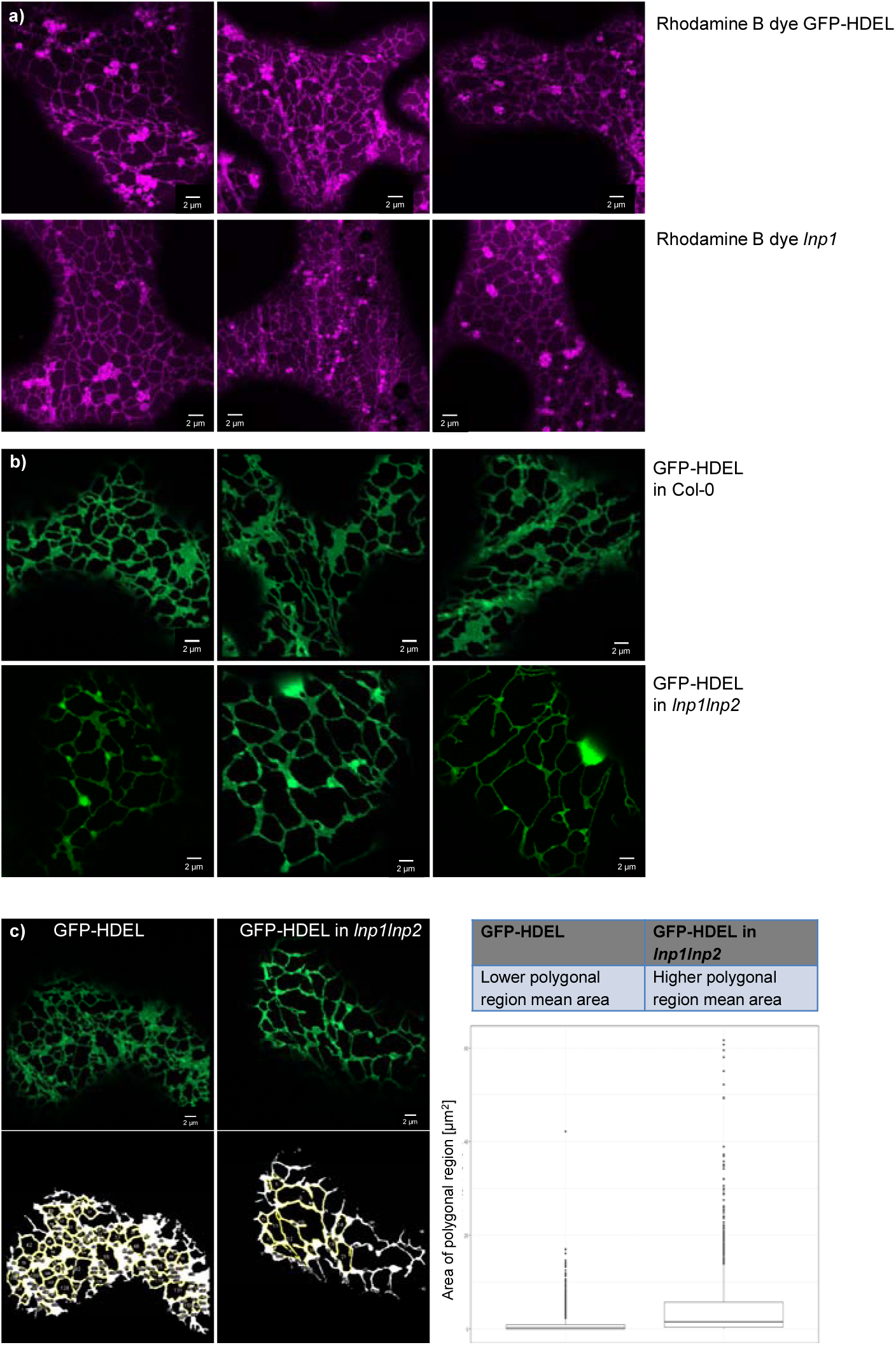
ER network structure in an arabidopsis lnp1 and lnp1lnp2 amiRNA mutant. a) The ER network in cotyledons of GFP-HDEL arabidopsis plants and a stable homozygous *lnp1* amiRNA line was visualised with the lipid dye Rhodamine B hexyl ester. This dye labels the ER network but also mitochondria. b) The ER network in cotyledons of Col-0 and stable homozygous *lnp1lnp2* amiRNA lines were visualised by transformation with the ER luminal marker GFP-HDEL. Representative images for GFP-HDEL in the wildtype Col-0 and in the *lnp1lnp2* amiRNA plants are shown. c) The areas of the polygonal regions in the ER network are outlined by GFP-HDEL in the wildtype Col-0 and in the *lnp1lnp2* amiRNA plants. Polygonal areas were quantified using ImageJ’s ‘Analyse particles’. The original confocal images are shown together with images post-processing. The yellow outlines show polygonal regions with their quantifications in the box plot below. Since the data is not normal the Wilcoxon rank sum test was applied. p-value < 2.2^e-16^ for three biological replicates each with at least 12 technical replicates.

### Interaction between LNP and RTN proteins

The ER morphology phenotypes observed upon overexpression or downregulation of AtLNP1 and/or AtLNP2 strongly suggest that LNP proteins are involved in the formation and/or stabilisation of trivia and cisternae in the ER network. Another class of proteins known to be required for the formation of a dynamic ER tubular network in plants are the RTN proteins which induce and/or stabilise membrane curvature and are capable of constricting tubules whilst suppressing cisternae formation (Sparkes *et al.*, 2010). Hence, we explored the interplay between RTN and LNP proteins.

Initially, the formation of protein-protein interactions between RTN1 and both LNP proteins were tested using Förster resonance energy transfer by fluorescence lifetime imaging microscopy (FRET-FLIM) analysis *in vivo* (Figure 5, Table 1, Supplementary Figure S5). RTN1 was selected as an exemplar of all RTN proteins since it has high sequence homology to other family members and is known to be expressed in all tissues throughout development (Arabidopsis eFP Browser; Winter *et al.*, 2007).

**Figure 5:**
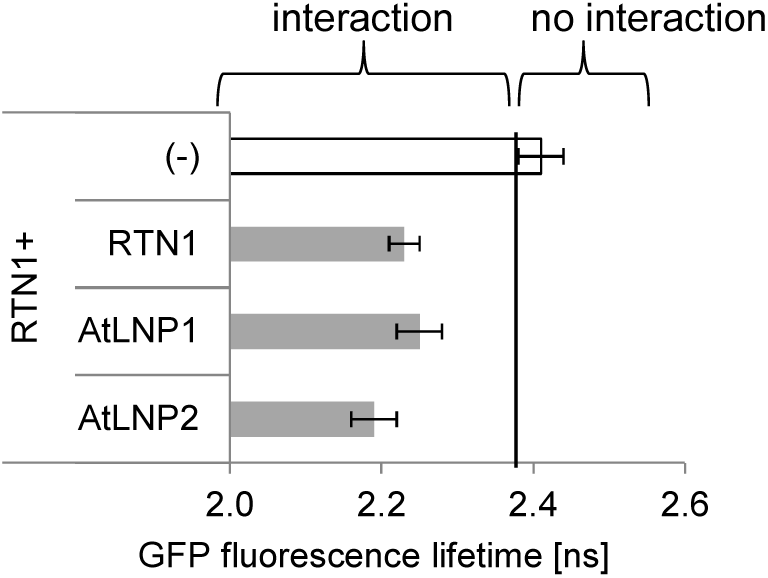
AtLNP1 and AtLNP2 form protein-protein interactions with RTN1. Combinations of donor (GFP-RTN1) and acceptor constructs (RFP-RTN1, AtLNP1-RFP or AtLNP2-RFP) were co-infiltrated into *N. tabacum* leaf epidermal cells, and protein-protein interactions assessed by FRET-FLIM analysis. For each measurement, a region of low-mobility ER continuous with the nuclear envelope was selected, and the fluorescence lifetime of the donor fluorophore was measured. Bar graphs depict the mean fluorescence lifetime (ns) ± SD. For each combination, at least two biological samples with a minimum of three technical replicates were used for the statistical analysis. The fluorescence lifetime of the donor construct (GFP-RTN1) in the absence of an acceptor was used as a negative control (white bar). Since RTN1 is known to form homo-oligomers, the fluorescence lifetime of GFP-RTN1 in the presence of the RFP-RTN1 acceptor was used as a positive control. Lifetimes significantly lower than those of RTN1-GFP alone (left side of the black line) indicate protein-protein interactions.

**Table 1:** Protein-protein interactions measured by FRET-FLIM. Donor and acceptor protein constructs are indicated together with the average fluorescence lifetime (in ns) for the donor fluorophore and the SD for each combination. For each combination, at least two biological samples with a minimum of three technical replicates were used for the statistical analysis.

| Donor (GFP) | Acceptor (mRFP) | Average [ns] | SE | $\Delta$ [ns] |
| --- | --- | --- | --- | --- |
| RTN1 | (-) | 2.42 | 0.03 | 0.00 |
|  | RTN1 | 2.21 | 0.03 | 0.20 |
|  | AtLNP1 | 2.20 | 0.02 | 0.21 |
|  | AtLNP2 | 2.20 | 0.03 | 0.20 |

Time-resolved fluorescence spectroscopy in imaging biological systems allows for the implementation of fluorescence lifetime imaging microscopy (FLIM). FRET-FLIM measures the reduction in the excited-state lifetime of GFP (donor) fluorescence in the presence of an acceptor fluorophore (e.g. mRFP) that is independent of the problems associated with steady-state intensity measurements. Reduction in GFP life-time is an indication that the two proteins are within a distance of 1 to 10 nm, thus indicating a direct physical interaction between the two protein fusions (Sparkes *et al.*, 2010; Schoberer and Botchway, 2014). It was shown previously that a reduction of as little as approximately 200 ps in the excited-state lifetime of the GFP-labelled protein represents quenching through a protein-protein interaction (Stubbs *et al.*, 2005).

Donor and acceptor constructs were co-infiltrated into *N. tabacum* leaf epidermal cells and FRET-FLIM analysis performed after 48 h to assess protein-protein interactions. For both AtLNP1 and AtLNP2, a significant reduction of 0.2 ns in the lifetime of the donor (LNP-GFP) fluorescence in the presence of the acceptor fluorophore (RFP-RTN1) was observed, in comparison to expression of the donor alone (Figure 5, Table 1, Supplementary Figure S5). These data indicate that AtLNP1 and AtLNP2 are both capable of physically interacting with RTN proteins *in vivo*.

Since FRET-FLIM analysis demonstrated that LNP and RTN1 proteins interact *in vivo*, the existence of a possible synergistic relationship between the two proteins which influences ER morphogenesis was evaluated *in planta*. AtLNP1 and AtLNP2 constructs were expressed using an OD_600_ of 0.3 that usually results in the formation of enlarged cisternae (Figure 3), RTN1 was infiltrated at a standard OD_600_ of 0.1 known to result in ER tubule restriction (Sparkes *et al.*, 2010). Transient coexpression of AtLNP1-GFP with RFP-RTN1 (Figure 6) appeared to suppress the proliferation of AtLNP1-labelled cisternae previously observed from infiltration of AtLNP1-GFP alone (Figure 6, compare with Figure 3). Instead AtLNP1-GFP labelling of trivia as well as smaller, partially-fragmented cisternae was detected. As observed when AtLNP2-GFP was infiltrated on its own (Figure 3), coexpression of AtLNP2-GFP with RFP-RTN1 resulted in the presence of both AtLNP2-containing sheets and tubules. In addition, overexpression of AtLNP2 together with RTN1 induced the formation of nodule-like structures in the ER tubular network (Figure 6).

**Figure 6:**
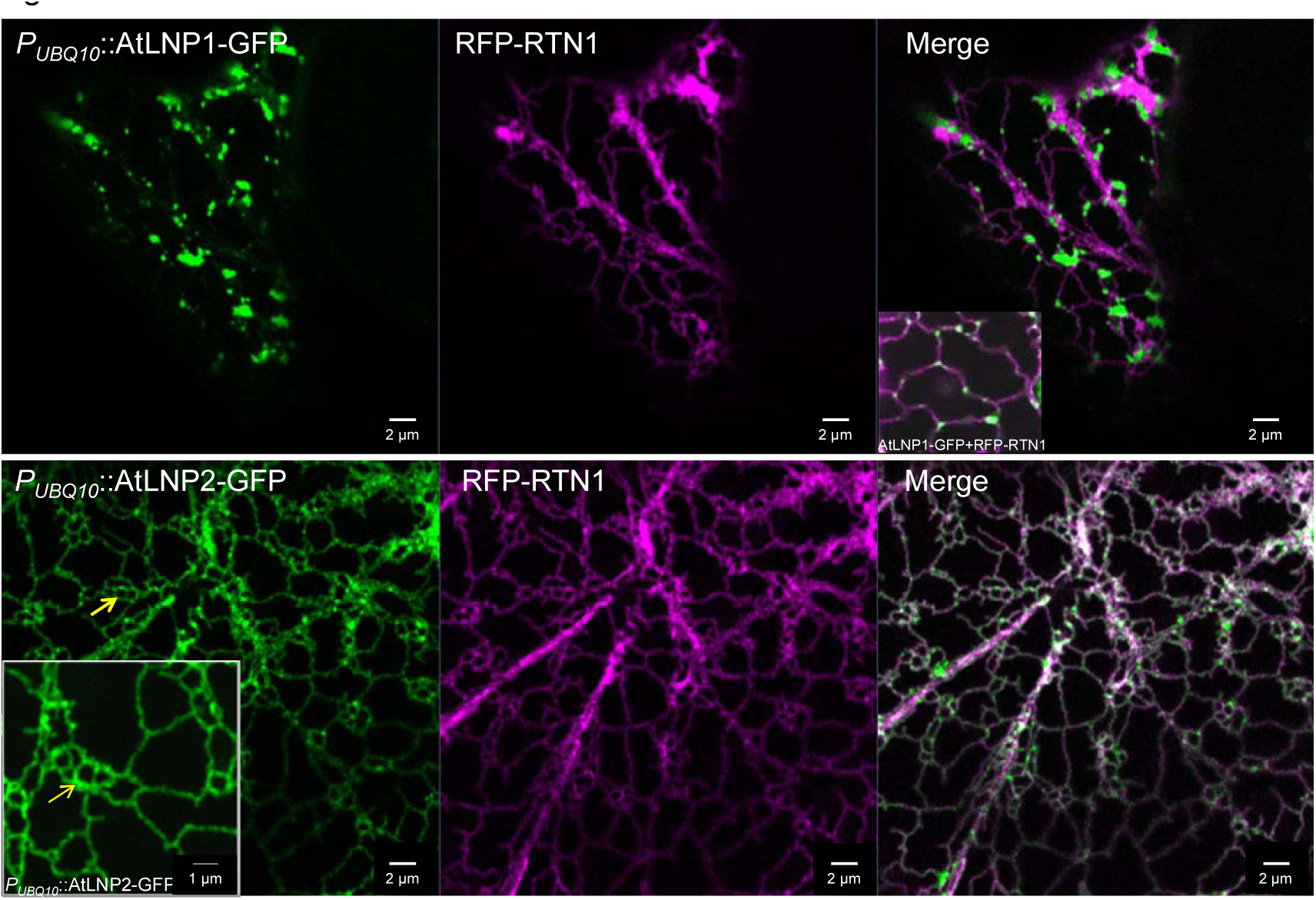
Overexpression of RTN1 influences the localisation of both AtLNP1 and AtLNP2. AtLNP1-GFP (*P*_*UBQ10*_∷AtLNP1-GFP, OD=0.3) and AtLNP2-GFP (*P*_*UBQ10*_∷AtLNP2-GFP, OD=0.3) were agroinfiltrated into *N. tabacum* leaf epidermal cells alongside RFP-RTN1 (35S∷RFP-RTN1, OD=0.1). Transient coexpression of AtLNP1 together with RTN1 results in the loss of large AtLNP1-labelled cisternae usually seen at this OD. Coexpression of RTN1 and AtLNP2, which is less sheet-specific and also localises to ER tubules, results in the formation of ER nodules (yellow arrow). Insets show more zoomed in images.

These data suggest that RTN1 is capable of counteracting the sheet-induction upon LNP overexpression, further strengthening the case for these proteins having a cooperative role *in vivo*.

### Dynamics of Lunapark-labelled cisternae and trivia

Yeast and mammalian Lnp proteins preferentially localise to the trivia of the ER and it has been suggested that in these organisms Lnp proteins are involved in stabilising these intersections (Chen *et al.*,: 2012; Chen *et al.*, 2015; Wang *et al.*, 2016). As AtLNP1 also preferentially localises to ER trivia we wanted to determine if AtLNP1 has a comparable function in stabilising the overall ER network.

The movement and remodelling dynamics of AtLNP1-labelled ER cisternae and trivia was investigated through time-lapse image processing. In general, several different types of motion were observed for arabidopsis LNP proteins that can be classified in similar categories as previously described for the mammalian Lnp1 (Chen *et al.*, 2015). These movements consist of stationary Brownian-like behaviour; directed movement of labelled regions along a tubule; merging of two adjacent puncta into a single punctum, and absorption and separation (mainly for cisternae) of discrete puncta into two independent puncta (Figure 7).

**Figure 7:**
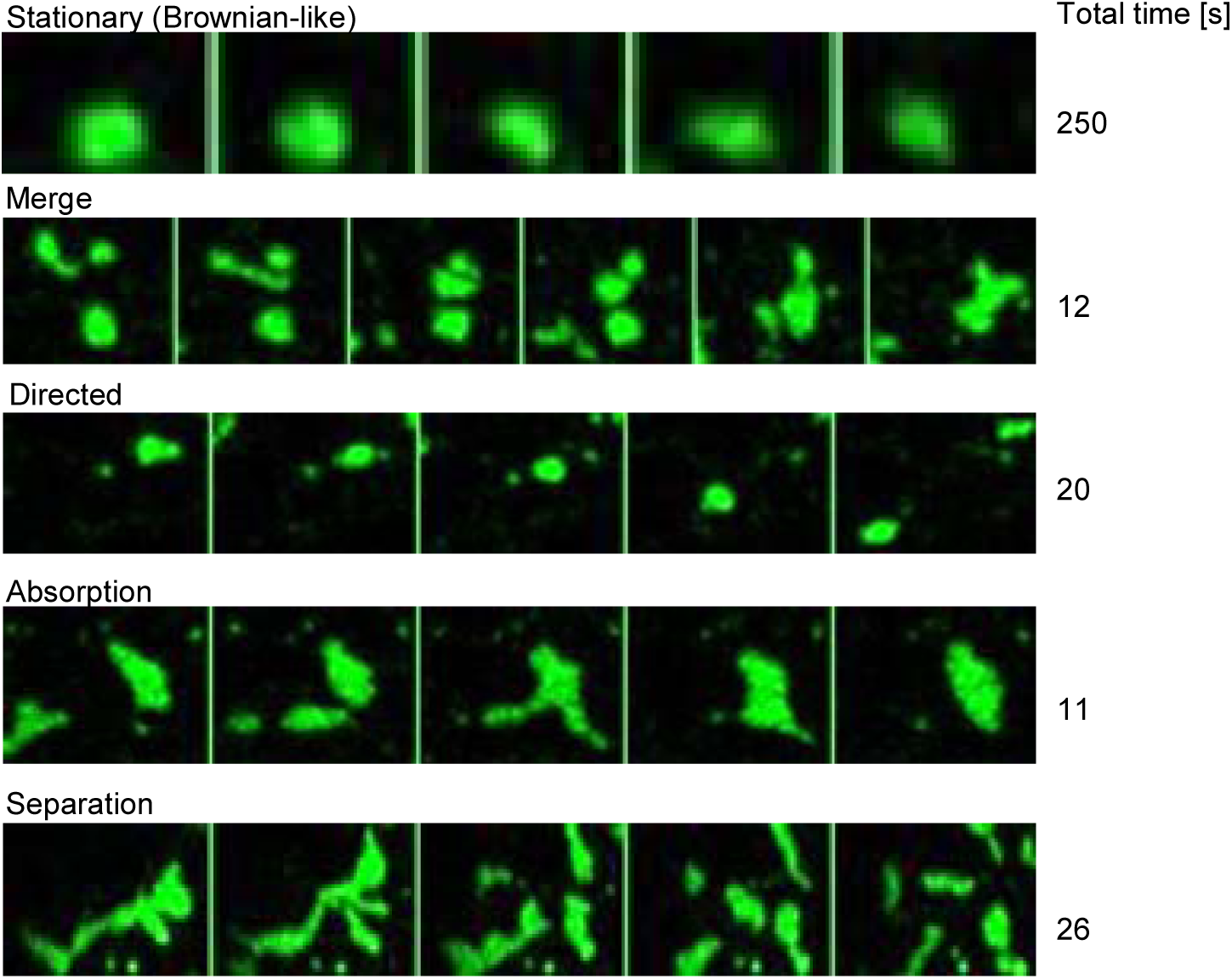
AtLNP1-labelled cisternae display different dynamic behaviours. Different types of movement dynamics observed in cisternae labelled with *P*_*UBQ10*_∷AtLNP1-GFP are listed by representative movies. The total time in seconds [s] of the movie shown is indicated on the right-hand side.

Further analysis of movies compiled from the time-lapse images (total of 30 images) revealed that out of the 90 AtLNP1-labelled puncta analysed in 20 movies, 59 (66%) moved from their original position. The remaining 31 (34%) puncta stayed fixed in their location and did not move. Moreover, within the duration of the time lapse experiment, 16 of these 90 AtLNP1-labelled puncta also fused together, whilst 7 puncta were absorbed into ER sheets. This appears to differ from the mammalian system cells where a significant majority (74%) of stable junctions are labelled with mLnp1 and less than 6% of unstable junctions actually acquire mLnp1 (Chen *et al.*, 2015).

To test if the presence of AtLNP1 actively stabilises the ER network or if AtLNP1-labelled junctions and cisternae follow the overall movement of the network as a whole, the dynamics of AtLNP1-labelled junctions and cisternae were analysed in comparison to the surrounding network (Figure 8). Persistency mapping was performed on movies capturing the movement of AtLNP1-GFP labelled trivia and cisternae (Figure 8a,b). Persistency was assessed using spatio-temporal projections of AtLNP1-GFP labelled cells over 60 s with analysis performed on both relatively stable (Figure 8c) and more dynamic (Figure 8d) regions of the cortical ER network, where the latter mainly represents *trans*-vacuolar strands. In the more stable areas of the ER where remodelling was minimal, AtLNP1-labelled ER elements similarly exhibited limited mobility and thus had high persistency (Figure 8b,c). It should also be noted that stable trivia and polygons were frequently observed to not have acquired AtLNP1. In areas of high motility such as vacuolar strands, AtLNP1-labelled areas also show high motility and low persistency (Figure 8b,d).

**Figure 8:**
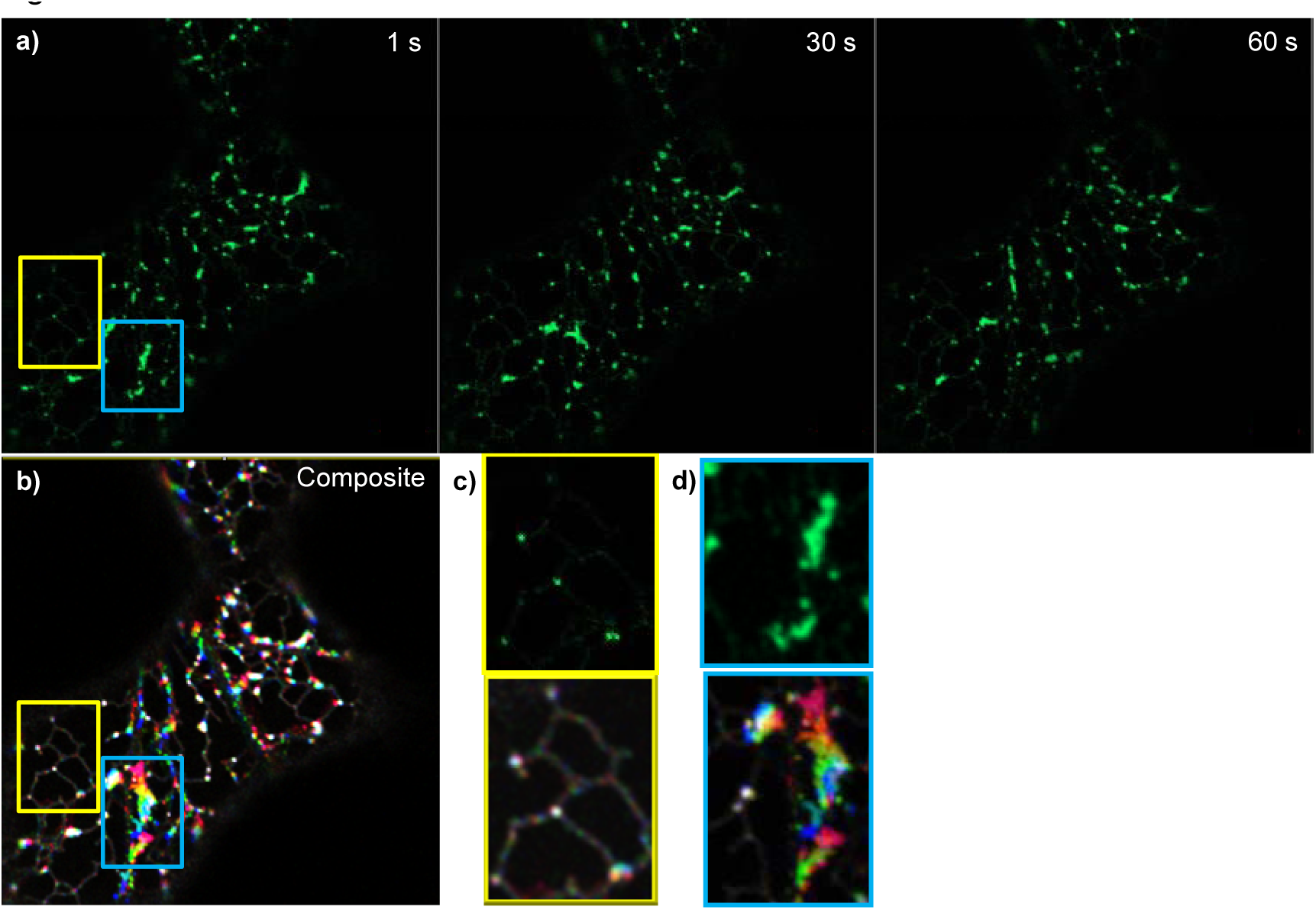
Persistency mapping of AtLNP1-labelled ER cisternae and three-way-junctions. a) Movies of ER cisternae labelled with P_UBQ10_∷AtLNP1-GFP were taken over 60 s. Static images of the AtLNP1-GFP-labelled ER network labelled are shown at 1 second, 30 seconds and 60 seconds. b) Composite image of the three timepoints shown in A following temporal colour coding of the movie (performed in the ImageJ ‘temporal color-code’ plugin). White areas indicate points of persistency occurring for the duration of the time imaged, coloured areas indicate movement with areas in red showing the highest mobility. c) Magnification of a stable region of the ER (framed in yellow in b). Confocal image of AtLNP1-GFP labelled trivia (top) and the corresponding persistency map (bottom). Within this relatively stable area of the ER network both AtLNP1-labelled junctions and junctions with no detectable AtLNP1 expression remain persistent. d) Magnification of a highly motile region of the ER undergoing rapid remodelling (framed in blue in b). Confocal image of AtLNP1-GFP labelled trivia and cisternae (top) and the corresponding persistency map (bottom).

Taken together, these data indicate that, unlike in mammalian systems, ER trivia and network stability is not dependent on the presence of AtLNP1 but that the motility of AtLNP1-labelled junctions and cisternae follows the motility or persistency of the surrounding ER network architecture structure.

## Discussion

Here we have reported, for the first time, the identification and characterisation of two plant homologues (AtLNP1 and AtLNP2) of the mammalian and yeast LNP proteins.

### Subcellular location of arabidopsis Lunapark proteins

AtLNP1 localises to trivia (three-way junctions) in the ER network. This subcellular localisation is comparable to that described for yeast and mammalian Lunapark proteins (Chen *et al.*, 2012; Shemesh *et al.*, 2014; Wang *et al.*, 2016). Both AtLNP1 and its animal counterpart can also label ER tubules at higher protein expression levels (Shemesh *et al.*, 2014; Wang *et al.*, 2016). In contrast, AtLNP2 expression was observed throughout the ER network, in both sheets and tubules, suggesting that, despite sharing 67% protein sequence homology with AtLNP1, they are not complete functional homologs. However, the lack of an obvious ER morphology phenotype in single loss-of-function AtLNP mutants (in contrast to that observed for the double *lnp1lnp2* mutant) would indicate some degree of functional redundancy. Furthermore, to date, no second Lunapark protein with an ER tubular localisation has been described in yeast and mammalian systems that would correspond with AtLNP2 raising the question about function and redundancy of the second arabidopsis Lunapark protein. *Homo sapiens*, for example, codes for at least five Lnp isoforms, which may potentially have different spatio-temporal expression patterns and/or functional specialisation.

### Lunapark function in the arabidopsis ER network

Overexpression of arabidopsis LNP proteins results in increased cisternae formation in a dose-dependent manner (Figure 3). In contrast, in mammalian cells it has previously been demonstrated that increasing amounts of mLnp1 protein result initially in ER cisternae induction, followed by the generation of thicker ER tubules at higher mLnp1 expression levels (Wang *et al.*, 2016). In COS cells (Shemesh *et al.*, 2014) mLnp1 localises to trivia at low expression levels as well as to some punctae on tubules. Higher mLnp1 expression levels result in clustered localisation to densely branched tubules also resulting in increased numbers of trivia; at very high expression levels Lnp localises to longer, unbranched tubules concomitant with a decrease in trivia. No such decrease in trivia abundance or thicker ER tubules were observed upon increased LNP expression in either tobacco or arabidopsis (Figures 2 and 3) although this may be due to the limitations of achievable maximal protein expression levels in the transient expression system since very high doses of agrobacterium frequently result in leaf necrosis.

Analysis of both loss and gain-of function mutants in the two identified AtLNP proteins revealed further potential functional differences between plant and mammalian/yeast Lunapark proteins. Knockdown of AtLNP1 alone does not result in a visible ER network morphology phenotype, however, a *lnp1lnp2* x GFP-HDEL knockdown line exhibited a significantly increased mean ER polygonal area in comparison to the wildtype x GFP-HDEL control (Figure 4). The polygonal areas of the mutant also spanned a greater size distribution than those of the wildtype control (Figure 4) ultimately resulting in a less structured, ‘looser’ ER network in the mutant. This observed ER morphology upon depletion of AtLNP again contrasts with that described in similar studies in yeast and mammalian systems. Mutations in the zinc finger motif of yeast Lnp1p led to a reduction in polygon size and thus resulted in a densely reticulated network (Chen *et al.*, 2012), and, similarly, the loss of mLnp1 gave rise to a more compact, sheet-like ER morphology (Chen *et al.*, 2015). LNP mutant forms expressed in U2OS cells lacking LNP show sheet generation with a reduction in tubules and junctions (Wang *et al.*, 2016). Moreover, addition of cytoplasmic fragments of *Xenopus* Lnp acting as a dominant-negative mutant to a *Xenopus* network formation resulted in the replacement of trivia by small cisternae as well as an overall reduction in trivia (Wang *et al.*, 2016). These results recently led Wang *et al.* (2016) to conclude that mammalian LNP is not essential for ER tubule and junction formation but instead affects trivia abundance.

The striking changes in polygonal structure and areas reported here for the arabidopsis *lnp1lnp2* knock-down line could also be due to a reduction in nascent trivia formation and/or loss of stability of trivia resulting in more irregular polygonal areas. However, it should be noted that we do not observe an increase in junctions (as described for yeast), or cisternae, as described in yeast and mammalian cells, respectively. An alternative explanation for the ER morphology observed in the *lnp1lnp2* mutant is that depletion of AtLNP1 and AtLNP2 may result in a decrease in cisternae and/or reduction in cisternal stability, which might also result in a change in the biophysical properties of the ER leading to a less stable and structured network. This hypothesis is supported by the formation of enlarged cisternal structures upon overexpression of arabidopsis Lunapark proteins, as described above (Figure 3).

Several functions for Lunapark proteins have been suggested in the various systems studied and are rather diverse. In *Caenorhabditis elegans* mutations in Inp-1 have been linked to neuronal defects similar to those in atlastin or the Yop1p homologue REEP1 mutants (Ghila and Gomez, 2008).

A function for mLnp1 in the stabilisation of trivia has been discussed (Chen *et al.*, 2015). Through immunolabelling of endogenous mLnp1, Chen *et al.* (2015) showed that the protein is only detectable in about half of the trivia in the mammalian network. The group also reported that junctions with mLnp1 are less mobile than junctions without mLnp1 and are less likely to show junction loss through ring closure (Chen *et al.*, 2015). A fraction of newly formed junctions go on to acquire mLnp1 protein. Newly formed junctions that do not acquire mLnp1 have a high probability of loss through ring closure and are relatively mobile, whereas conversely, those that do acquire mLnp1 have a greatly reduced probability of loss and are less mobile. This is especially prominent in newly formed junctions indicating that mLnp1 stabilizes newly formed trivia but is not required to be continually present on the junction there after (Chen *et al.*, 2015). In the absence of mLnp1, new junctions are still being formed but are less likely to persist resulting in more sheet-like structures (Chen *et al.*, 2015). For the plant system we could not find any indication that AtLNP1 or 2 stabilise trivia. The network dynamics is instead dependent on the surrounding network rather than the presence or absence of Lunapark proteins (Figure 8).

For mammalian cells it was also suggested that mLnp1 is not necessary for the generation or maintenance of the ER network since in the absence of mLnp1 there is still a reticular network, even if most trivia are converted into larger cisternae (Wang *et al.*, 2016). Instead, mLnp1 proteins are proposed to move into trivia with the overexpression of mLnp1 resulting in the expansion of cisternae but atlastin proteins are suggested to be responsible for the initial formation of native junctions (Wang *et al.*, 2016).

Interestingly, a theoretical model predicts mLnp1 to be a so-called S-type protein capable of stabilizing curvature and favouring negative curvature, which plays an important role in generating and stabilizing three-way tubular junctions (Shemesh *et al.*, 2014). This contradicts the hypothesis of Chen *et al.* (2012) that Lunapark proteins are involved in abolishing trivia in yeast. The latter was suggested following the observation that the ER in lnp1p mutants is highly reticulated resulting in an increased abundance of trivia. However, Shemesh *et al.* (2014) suggest that this might well be a cisternal structure rather than a reticulated network since the two cannot be distinguished at the resolution of the images. The model of Shemesh *et al.* (2014) is also of relevance to arabidopsis Lunapark proteins since cisternae are often bordered by negatively curved edge lines and it may hint at a mechanism for how AtLNP1 and 2 induce or stabilise cisternae.

### Interaction with reticulons

In yeast, it is suggested that Lnp1p acts in synergy with the reticulons and Yop1p. Indeed, Lnp1p has been shown to interact with Rtn1p, Yop1p and Sey1p since a loss of function mutation in Lnp1p in an rtn1/rtn2/yop1 triple mutant results in growth and ER morphological defects (Chen *et al.*, 2012). Lnp1p physically interacts with Rtn1p indicating that they may act on converging pathways since mutants in these genes display different ER phenotypes: loss of Lnp1p leads to the formation of densely reticulated ER (Chen *et al.*, 2012), whereas the loss of Rtn1p results in non-fenestrated sheets (DeCraene *et al.*, 2006).

In mammalian cells it has been proposed that Lnp proteins are not required for ER network formation but instead are involved in the formation of sheets at tubule junctions (Wang *et al.*, 2016). The presence of Lnp within the trivia may then prevent the migration of reticulons into the junction, thereby preventing junction expansion; in the absence of LNP trivia could therefore expand into large sheets (Wang *et al.*, 2016).

We show here that the AtLNP1 and 2 proteins are capable of interacting with reticulons (Figure 5) and, moreover, the induction of large cisternal regions resulting from LNP overexpression is suppressed by coexpression of reticulon proteins (Figure 6). We therefore hypothesise that, in arabidopsis, Lunapark and reticulon proteins function in concert to maintain the ratio of ER cisternae to tubules and are capable of interconverting the two morphologies.

In conclusion, we propose that arabidopsis Lunapark proteins are involved in the formation and/or stabilisation of ER network cisternae. They are also highly likely to functionally cooperate with other ER morphogens such as the reticulon family of proteins, with which they interact *in planta*. In contrast to that described for their yeast and mammalian orthologs, we did not find any evidence that arabidopsis Lunapark proteins stabilise ER trivia. The presence of two Lunapark homologs within the arabidopsis genome that show distinct subcellular localisation but also have some degree of functional redundancy, as shown in the requirement for a double knockdown, is of interest. We are currently investigating the impact of the different localisations of AtLNP1 and 2 on the ER and resulting functionality as well as the interactions of the AtLNP proteins with other ER-shaping proteins.

## Accession numbers

AtLNP1: At2g24330.1, AtLNP2: At4g31080.1, AtLNP2.2: At4g31080.2

## Acknowledgements

The authors thank Dr Imogen Sparkes (University of Bristol) for initial discussion on the Lunapark work and Samuel Williams for technical support. This work was supported by the British Biotechnology and Biological Sciences Research Council (grants BB/J017582/1 to LF and BB/M000168/1 to CH) via the ERA-CAPs programme “Per Aspera” (grant no. BB/J004987/1), the Science and Technology Facilities Council Program (grant no. 14230008 to CH), and a Vice Chancellor’s Fellowship from Oxford Brookes University to VK.

## Author contributions

V.K. and C.H. designed the research. E.B. and L.F. cloned the constructs, generated the mutant lines and performed the LNPARK amino acid comparative motif analysis. C.P. and F.T. carried out the mutant and persistency analysis. V.K. carried out confocal microscopy and the overexpression and FRET-FLIM analysis. All authors contributed to the writing of the article.

